## Supplementary Information for "Beam image-shift accelerated data acquisition for near-atomic resolution single-particle cryo-electron tomography"

**Supplementary Table 1**. Data collection statistics for tilt-series acquired using BISECT. Monodisperse (SPA) and pleomorphic (CET) samples with varying ice-thickness were acquired using different tomographic imaging conditions. All datasets were collected using a 5x5 BIS pattern. The number of ROIs imaged per BIS area for the pleomorphic samples was lower than the monodisperse samples because we skipped empty holes (the average number of imaged ROIs is reported in this case). CTF-fit resolution values for each session are reported as indicators of optical image quality (max/mean CTF-fit values were calculated over all tilted images in a dataset).

| **Sample Name/Type** | dNTPase | | Monodisperse | | Pleomorphic | |
|---|---|---|---|---|---|---|
| **Ice thickness (nm)** | <100 | | <100 | | 100-300 | |
| **Camera** | K2 | K2 | K3 | K3 | K2 | K3 |
| **Pixel Size (Å)** | 1.37 | 1.37 | 1.69 | 1.38 | 1.37 | 1.37 |
| **Tilt-Range ($^o$)** | ±60 | ±36 | ±60 | ±36 | ±60 | ±36 |
| **Number of Tilts** | 41 | 25 | 41 | 25 | 41 | 25 |
| **Total Dose ($e^-$/Å$^2$)** | 120 | 120 | 120 | 120 | 120 | 120 |
| **Areas Imaged** | 5 | 11 | 9 | 13 | 44 | 62 |
| **ROIs per BIS Area** | 25 | 25 | 25 | 25 | 10 | 16.4 |
| **ROIs per Hole** | 1 | 1 | 1 | 1 | 1 | 4 |
| **Number of Tilt-Series** | 125 | 275 | 225 | 325 | 444 | 1017 |
| **Max/Median CTF-fit Resolution (Å)** | 3.1/7.2 | 3.0/5.3 | 3.1/6.7 | 3.2/6.8 | 3.1/10.7 | 3.1/5.2 |
| **Total Time Per Tilt-Series (min)** | 7.6 | 4.7 | 3.0 | 2.1 | 14.7 | 2.4 |

**Supplementary Table 2**. Data processing statistics for CET and SPA datasets of dNTPase. All images were collected using a 5x5-hole lattice with the stage centered on the target area and each hole acquired using BIS. The two CET datasets were processed using SVA/CSPT and the single-particle dataset was subjected to SPA refinement.

| **Modality** | BISECT/CSPT | BISECT/CSPT | SPA |
|---|---|---|---|
| **Pixel Size (Å)** | 1.37 | 1.37 | 1.37 |
| **Number of Frames** | 4 per tilt (41 tilts per ROI, ±60º) | 11 per tilt (25 tilts per ROI, ±36º) | 60 per ROI (no tilting) |
| **ROIs Imaged** | 125 (64 selected) | 275 (64 selected) | 2,275 (64 selected) |
| **Number of Sub-Volumes/Particles** | 29,925 | 34,435 | 34,190 |
| **Estimated Resolution (Å)** | 4.9 | 3.6 | 3.3 |

**Supplementary Figure 1**. Implementation of workflow for tilt-series acquisition using BISECT and acquisition times per tilt-series as a function of the number of ROIs. **(A)** Middle branch shows the standard protocol for tilt-series acquisition. The box on the right shows the tracking steps, and the box on the left shows the targeting steps. Operations done by SerialEM and our custom Python routines are shown in purple and blue, respectively. **(B)** Curves showing total acquisition time per tilt-series as a function of the number of imaged ROIs using BISECT. Acquisition times measured for the K2 and K3 detectors are shown in blue and orange, respectively. Theoretical lowest times per tilt-series shown with dashed lines. Calculations were made based on experimental average times assuming 41 images per tilt-series, 3 $e^{-}/Å^{2}$ per tilt, and the same tracking accuracy for each ROI.

**Supplementary Figure 2**. Per-tilt astigmatic CTF estimation on tilt-series from different CET samples**.** Slices through tomographic reconstruction (top), fitted CTF model and corresponding 1D radial profile (red), estimated defocus (green dashed line), and CTF-fit scores (blue) for projections at high (51°), intermediate (24°) and low (0°) tilts (bottom), for tilt-series downloaded from EMPIAR-10304, EMPIAR-10453 and EMPIAR-10376, corresponding to *E. coli* 70S Ribosomes [32], intact SARS CoV-2 spike protein on intact virions [43], and lamella from yeast cells generated by FIB milling [37]. Scale bars are 300nm, 200nm and 700nm, respectively. Defocus values reported as [DF1, DF2, AST] are as follows: EMPIAR-10304 at 51° [22368, 21768, 5.15], at 24° [22035, 21670, -52.17], and at 0° [20694, 20496, -1.31]. EMPIAR-10453 at 51° [16924, 16724, -32.83], at 24° [17559, 17303, -39.15], and at 0° [17282, 17204, -23.18]. EMPIAR-10376 at 51° [96413, 95698, -47.99], at 24° [95816, 93952, 84.33], and at 0° [94617, 92942, 63.48]. Defocus values (DF1 and DF2) are expressed in Å and astigmatism angles (AST) in degrees.

**Supplementary Figure 3**. Derivation of per-particle defocus based on Z-height values**. (A)** Schematic for derivation of per-particle CTF parameters. Defocus at the position of the tilt-axis is calculated as: $h * cos\theta + x * tan\theta$, where $h$ is the distance of the specimen to the tilt-axis, $x$ is the distance of the specimen to the central point (where the defocus is actually measured), and $\theta$ is the tilt-angle. The height change for the per-particle correction is estimated from: $cos\theta * (P_z + P_x * tan\theta)$, where $P_x$ is the particle's distance to a plane through the tilt-axis and orthogonal to the specimen plane, and $P_z$ is the relative height of the particle (red dot) with respect to the plane of the tilt-axis. **(B)** XZ cross-section through tomographic reconstruction showing the relative height at the specimen position with respect to the position of the tilt-axis (blue dot).

**Supplementary Figure 4**. Resolution improvement obtained by CSPT on mammalian 80S ribosome dataset (EMPIAR-10064). **(A-C)** Overview of reconstruction (top), density for an RNA helix (middle), and alpha helical segments (bottom) with fitted atomic coordinates PDB ID 4V7R for the 8.4 Å reconstruction obtained using EMAN2 (EMD-0529) [17] **(A)**, the 5.7 Å map obtained using M (EMD-11654) [7] **(B)**, and our 5.6 Å resolution reconstruction **(C)**. **(D)** Corresponding FSC curves between half-maps with resolutions measured according to the 0.143-cutoff criteria.

**Supplementary Figure 5**. Resolution improvement obtained by CSPT on *E. coli* 70S Ribosome dataset (EMPIAR-10304/EMD-10211). **(A)** Overview of reconstruction (left), alpha helical segments (middle), and density for a segment of RNA (left) with fitted atomic coordinates PDB ID 5MDZ for the 7 Å resolution reconstruction obtained using emClarity [41]. **(B)** Corresponding panels for the 4.8 Å resolution reconstruction obtained using CSTP. **(C)** FSC curve between half-maps for the CSPT reconstruction showing the estimated resolution according to the 0.143-cutoff.

**Supplementary Figure 6**. Resolution improvement obtained by CSPT on RyR1 ion channel dataset (EMPIAR-10452/EMD-10840). **(A)** Overview of reconstruction (left), zoomed in view (middle), and density for alpha helical segments (left) with atomic coordinates PDB ID 5GKY fit into the original 9.1 Å resolution reconstruction obtained using dyn2rel [33]. **(B)** Corresponding panels for our 8.0 Å resolution reconstruction obtained using CSTP. **(C)** FSC curve between half-maps for the CSPT reconstruction.

**Supplementary Figure 7**. Distribution of positions selected for processing from the BIS pattern according to distance from the tilt-axis. Out of a total of 275 tilt-series collected over 11 areas, the best 64 tilt-series were selected for data processing based on the estimated CTF resolution. Histogram shows that selected positions were spread across a range of distances from the tilt-axis.

**Supplementary Figure 8**. Incremental improvement in resolution obtained for the dNTPase CET/SVA structure**. (A-B)** Half-map FSC curves **(A)** and density for an alpha helix with fitted atomic model **(B)** for CSPT reconstructions obtained at different stages of processing. Baseline reconstruction obtained using a single non-astigmatic defocus per tilt-series and no exposure weighting (5.8 Å, green), reconstruction obtained using a per-tilt CTF model without (5.0 Å, blue) and with astigmatism correction (4.6 Å, magenta), map obtained using theoretical exposure weighing as implemented in the IMOD package (4.3 Å, gray), and final map obtained using our astigmatic per-tilt CTF model and the self-tuning exposure weighting filter (3.6 Å, red).

**Supplementary Figure 9**. Comparison of CET and SPA structures of dNTPase**. (A)** Half-map FSC curves of reconstructions determined by BISECT/CSPT from tilt-series collected using tilt-ranges of ±60° (green) and ±36° (blue) with 3° increments, and SPA reconstruction obtained from micrographs acquired without tilting (red). Estimated resolutions according to the 0.143-criteria are 4.9 Å, 3.6 Å and 3.3 Å, respectively. **(B)** Comparison of secondary structure features between ±36° BISECT/CSPT and SPA maps (regions correspond to those presented in **Figure 5D)**.

**Supplementary Video 1**. Raw tilt-series of the tracking area obtained using BISECT. Sequence of 41 unaligned raw tilted projections collected over a ±60° tilt range acquired using BIS. We can achieve tracking with precision greater than 5 nm over the course of the tilt-series by using crosscorrelation maximal distance thresholds of 50 nm at the low-magnification level, and 2 nm at the high-magnification level. Field of view is ~500 nm.

**Supplementary Video 2**. Raw tilt-series of a ROI obtained using BISECT**.** Representative area situated 5.4 µm away in x along the tilt axis and 4.8 µm perpendicular to the tilt axis from the tracking area.

### Supplementary Figure 1

**A**

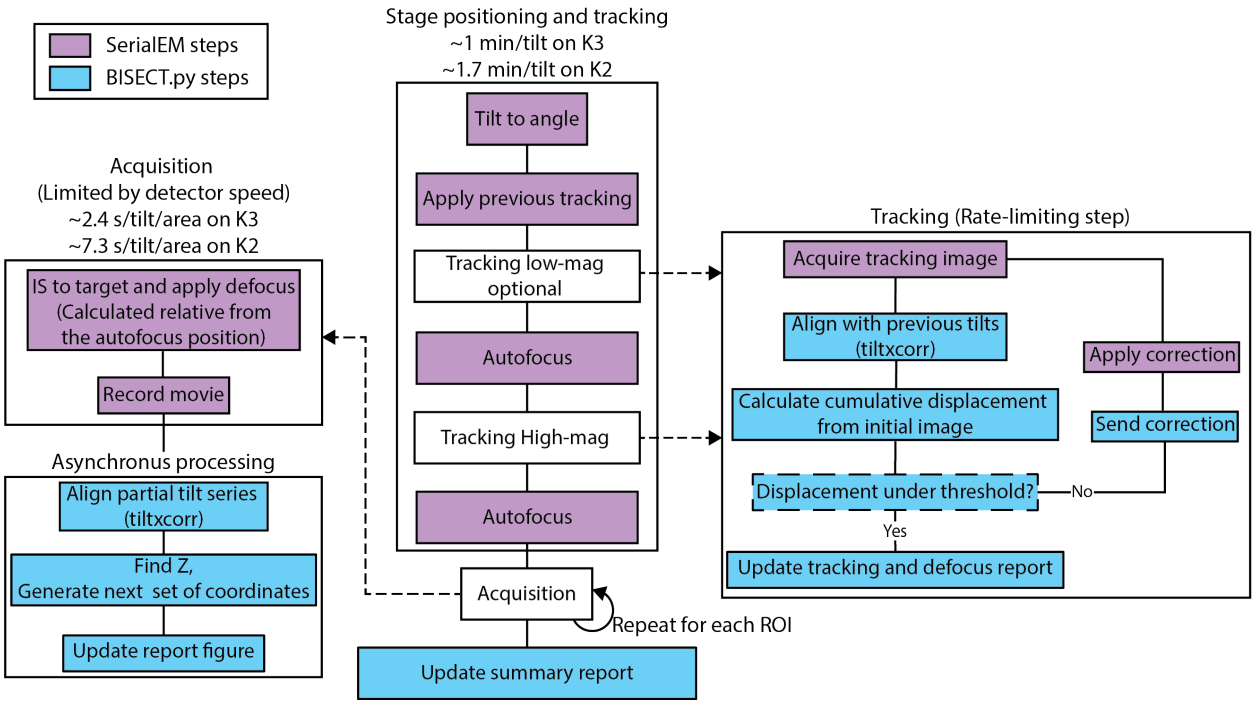


**B**

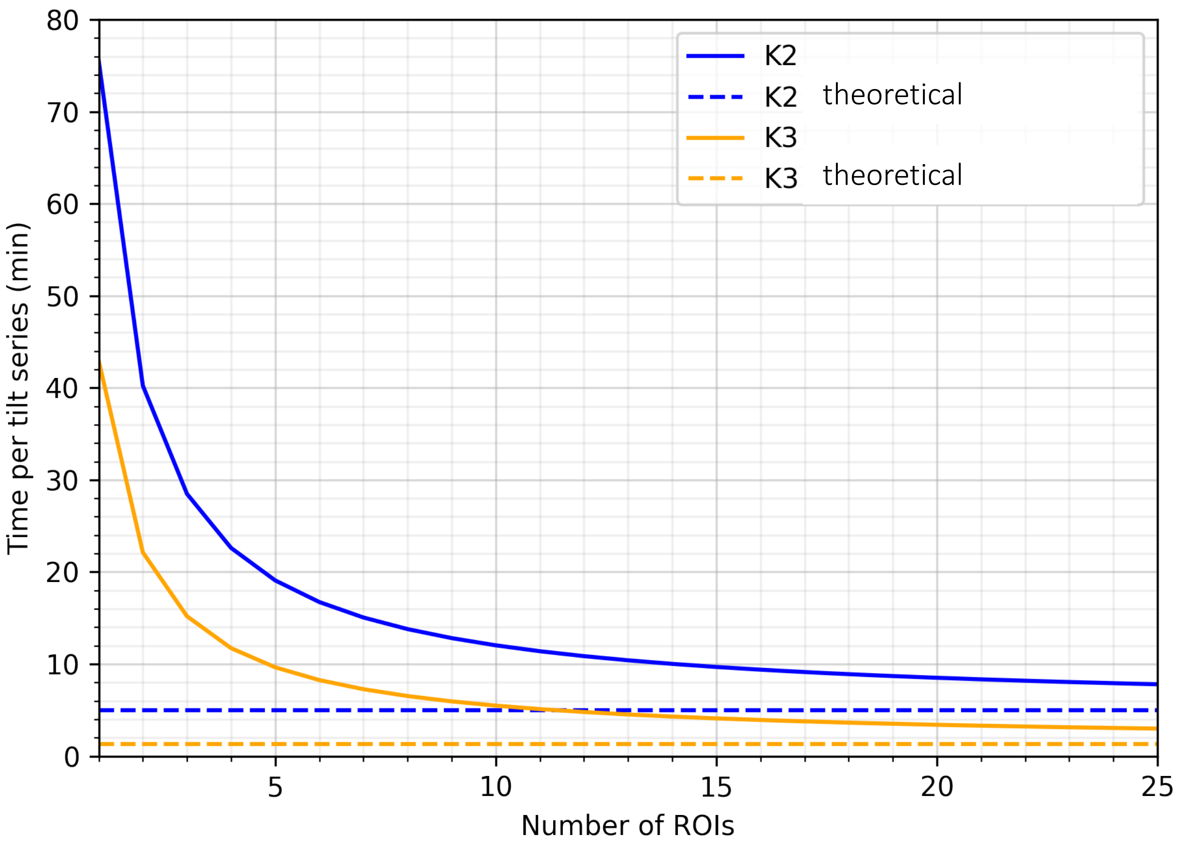

### Supplementary Figure 2

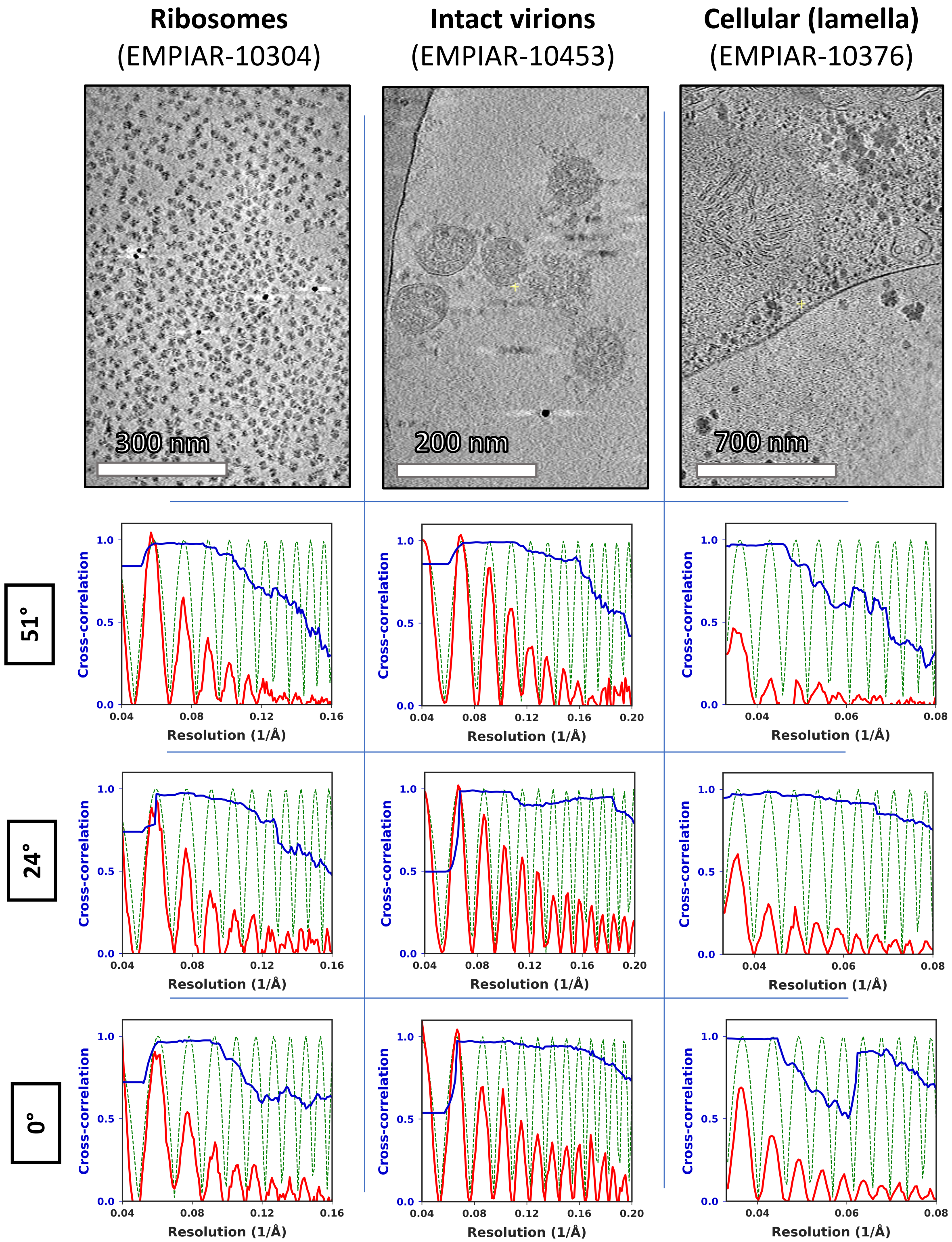

#### Supplementary Figure 3

**A**

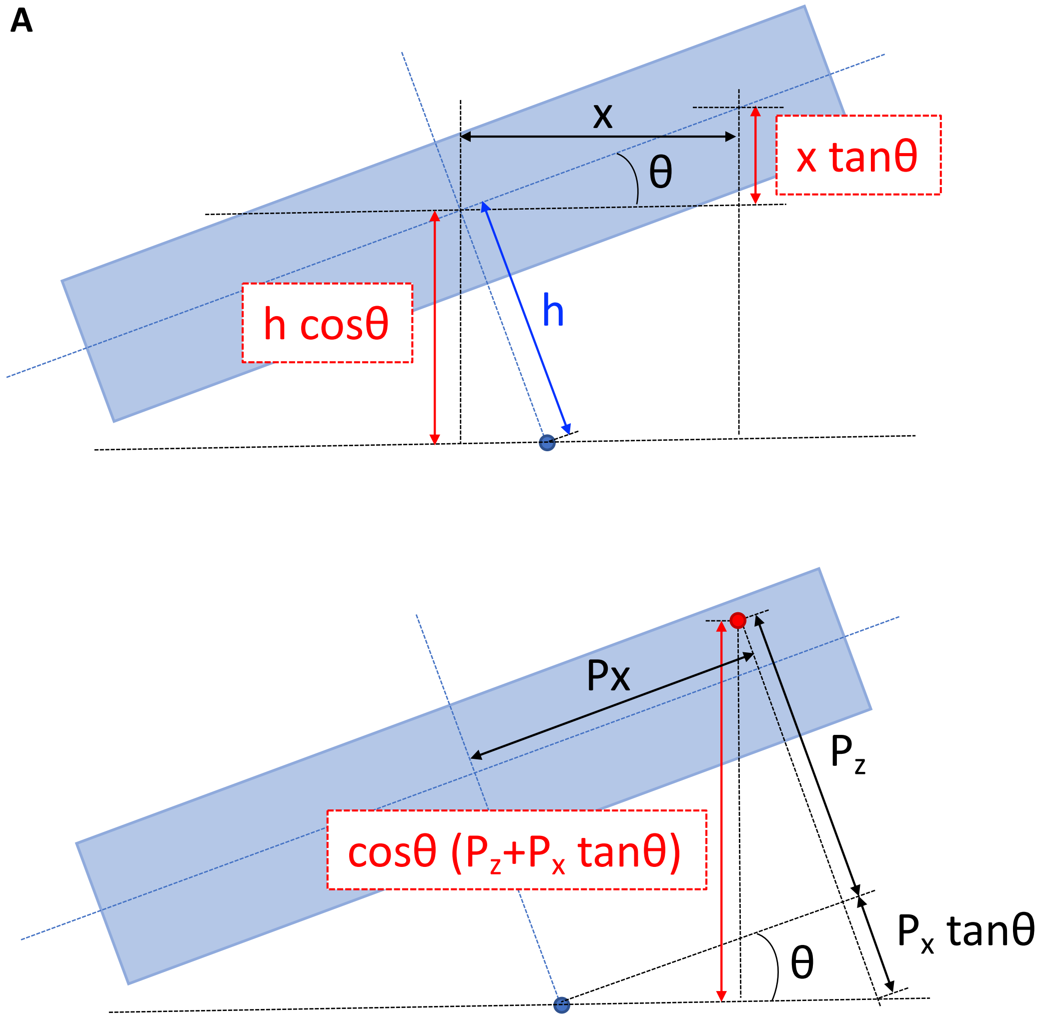


**B**

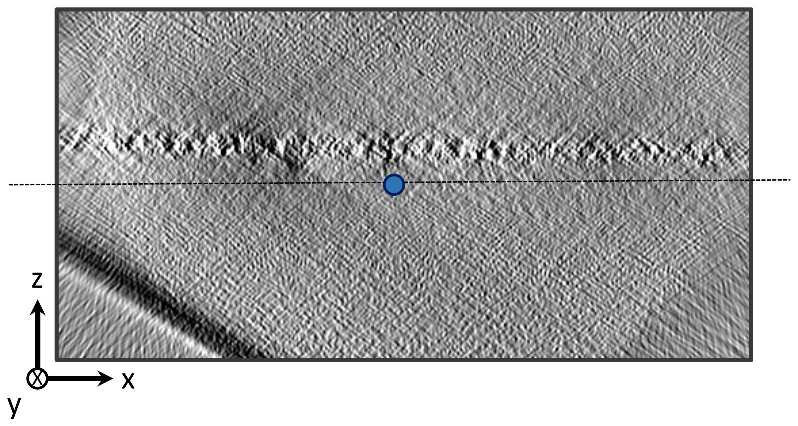

### Supplementary Figure 4

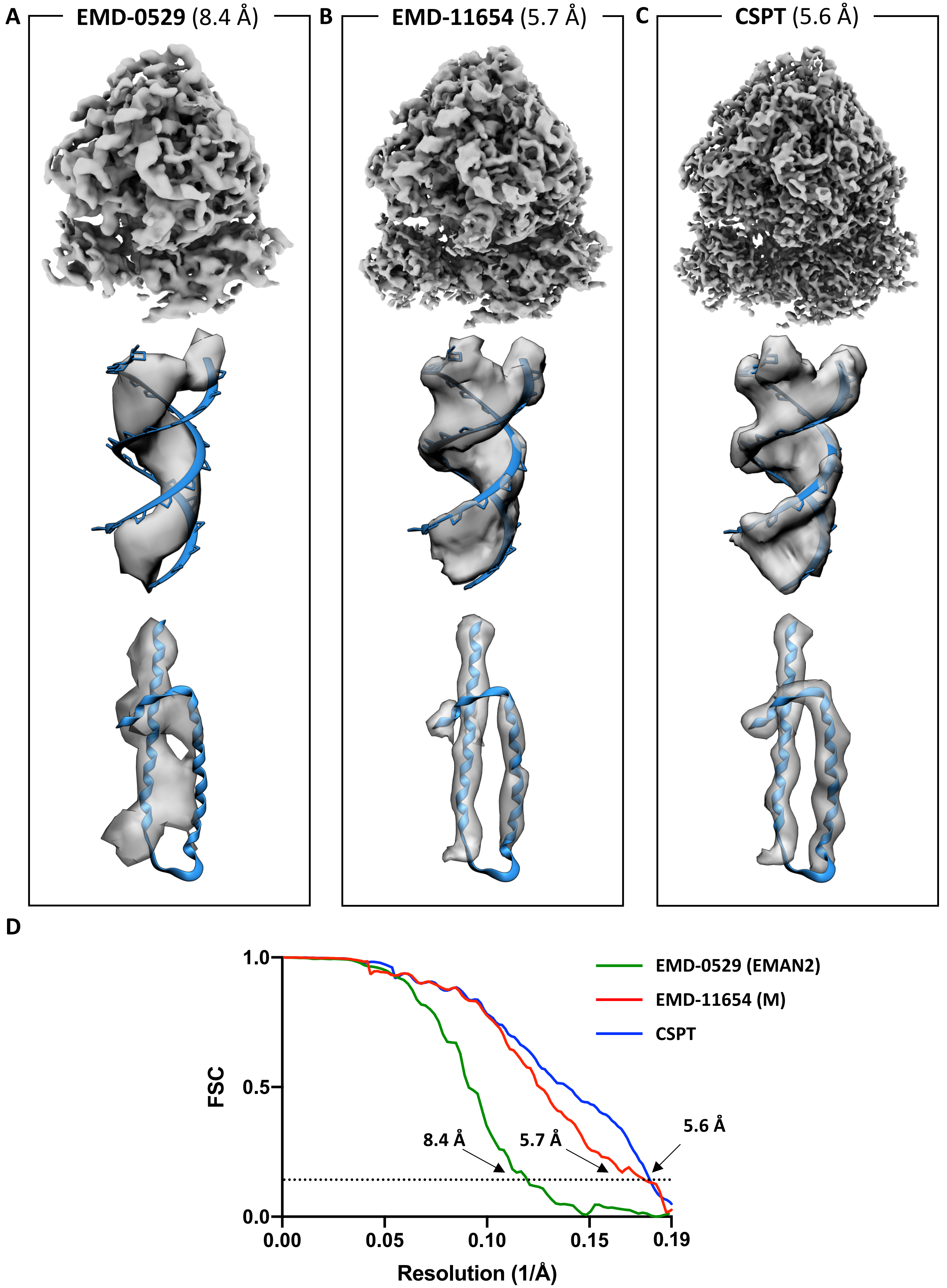

#### Supplementary Figure 5

A

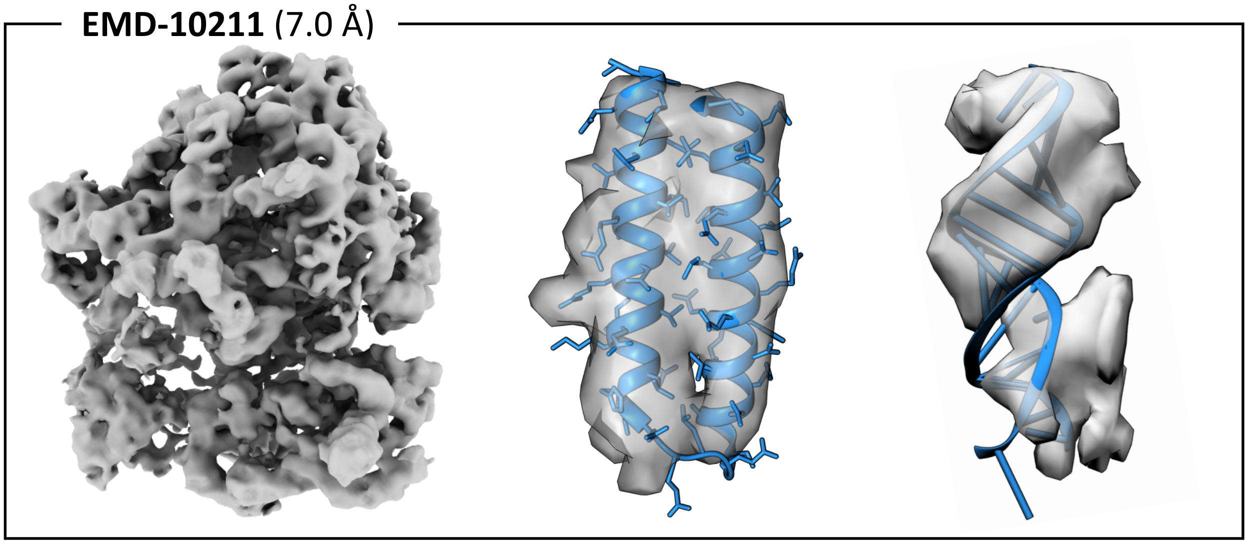


B

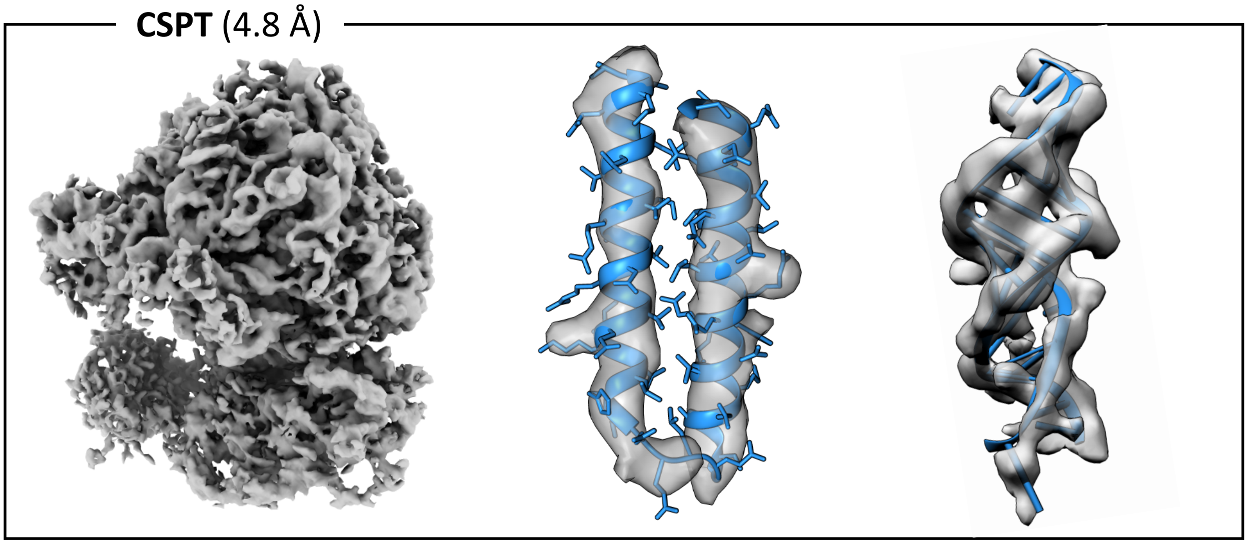


C

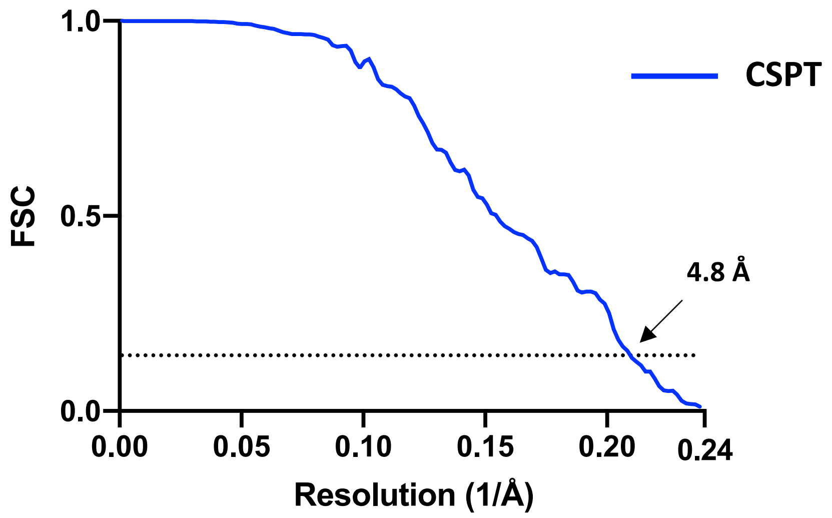

#### Supplementary Figure 6

A

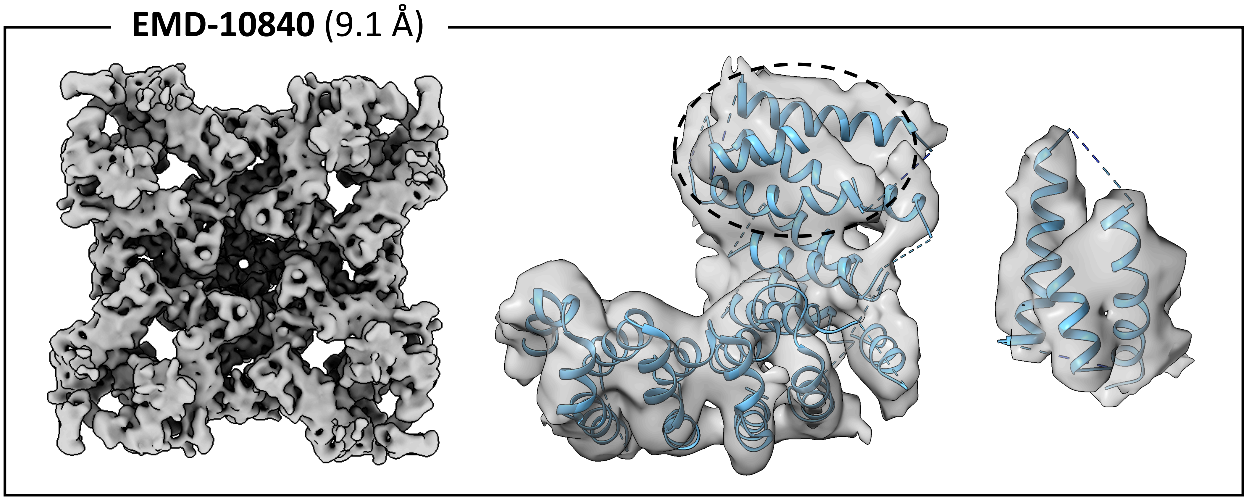


B

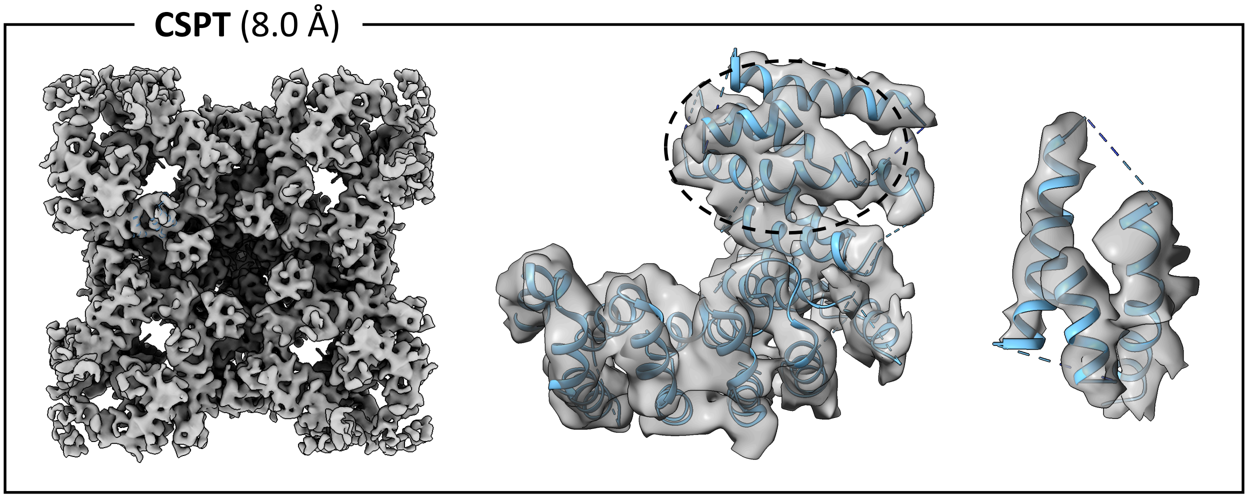


C

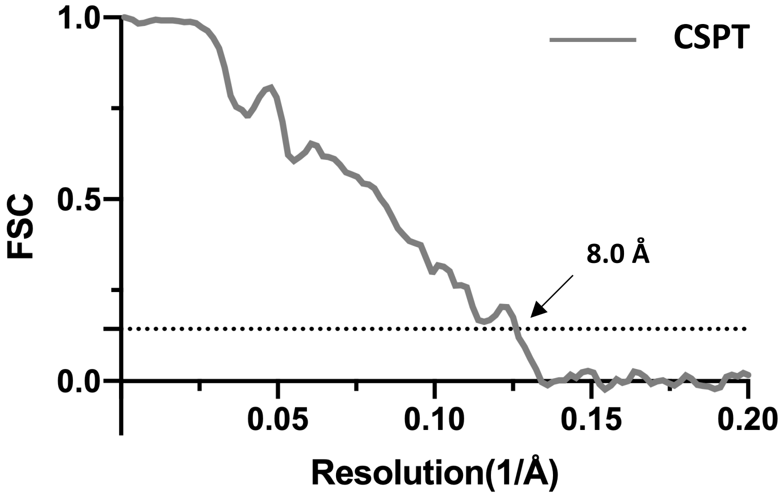

**Supplementary Figure 7**

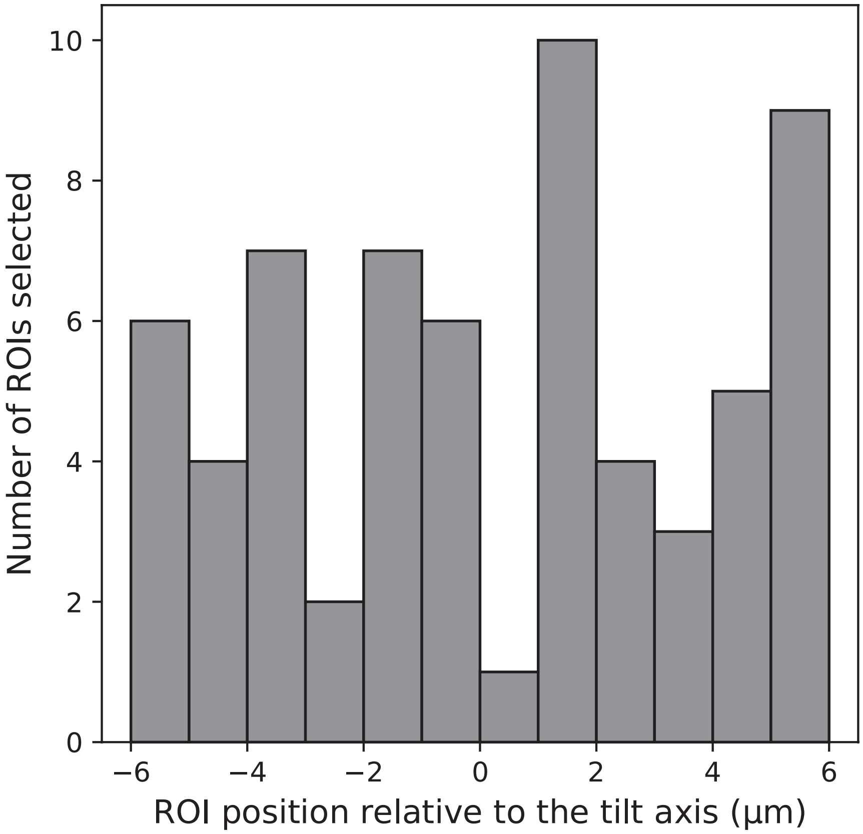

### Supplementary Figure 8

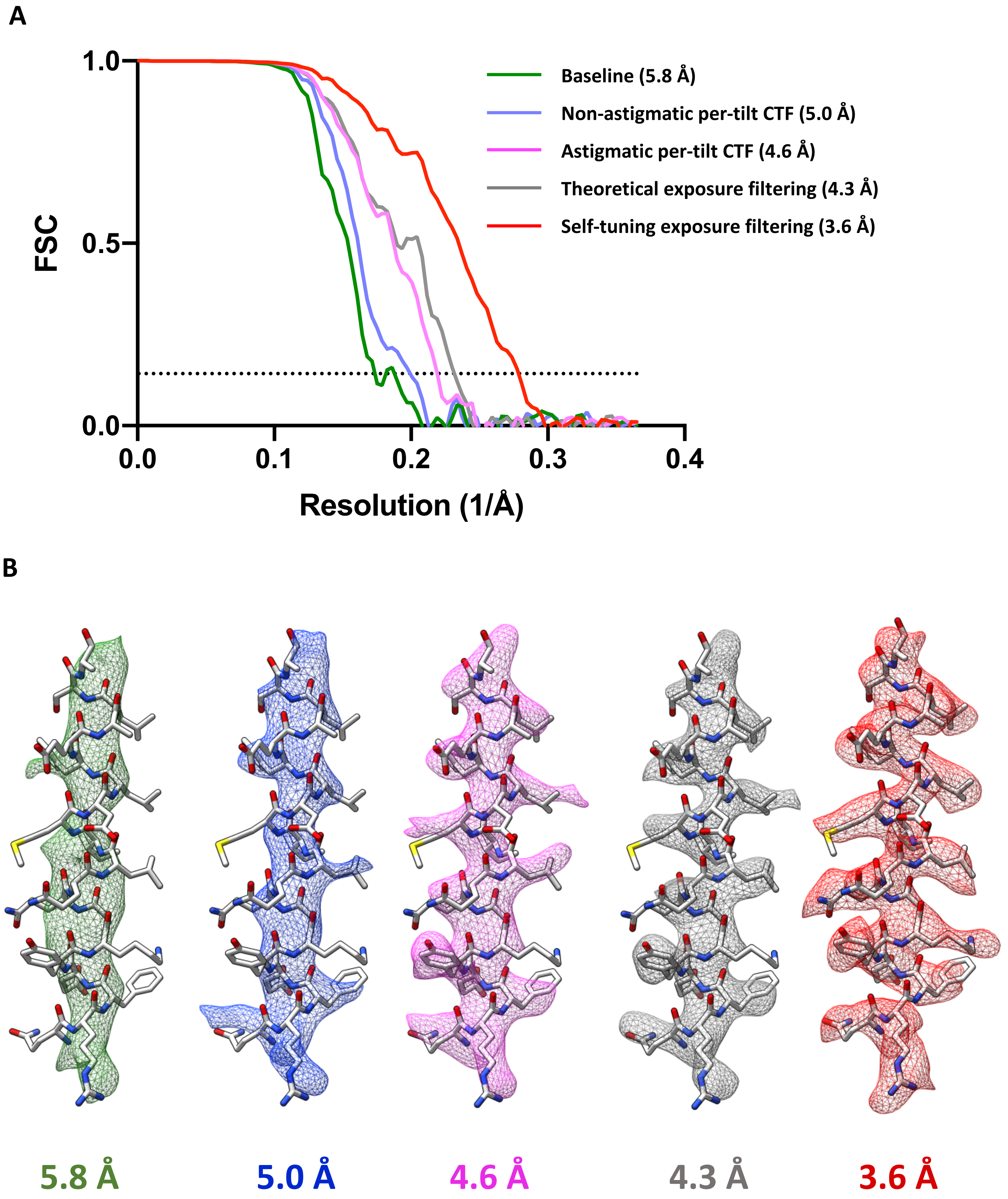

#### Supplementary Figure 9

A

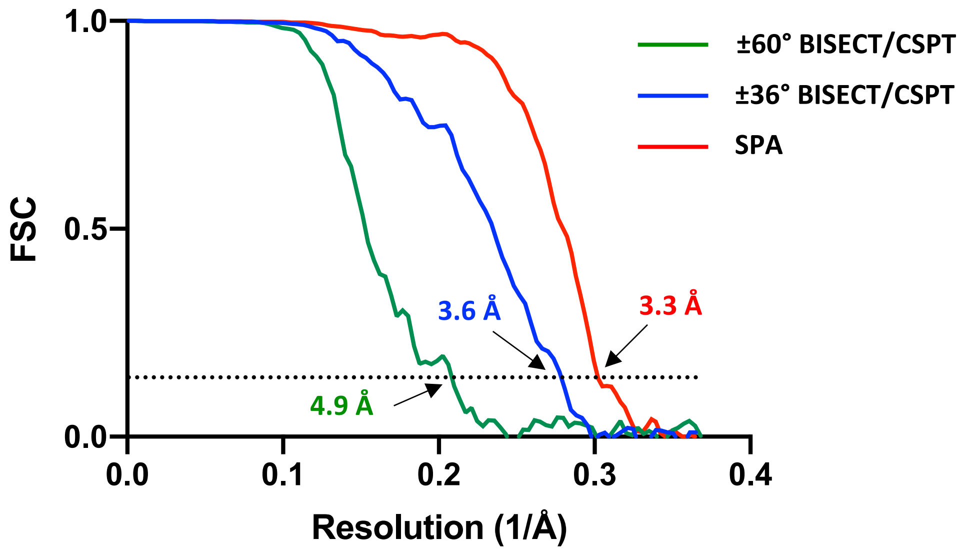


B

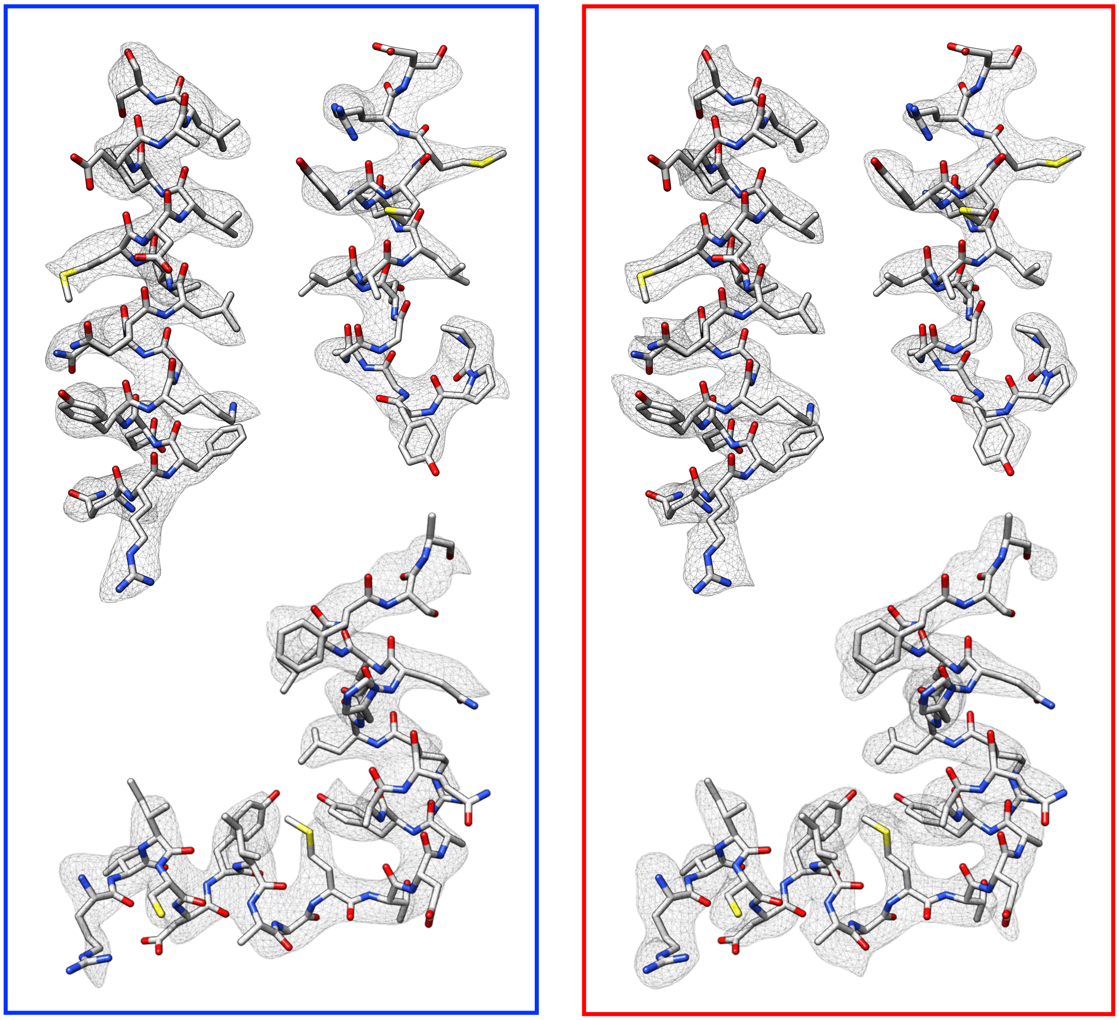

±36° BISECT/CSPT

SPA
